# Social effects of territorial neighbours on the timing of spring breeding in North American red squirrels

**DOI:** 10.1101/329276

**Authors:** David N. Fisher, Alastair J. Wilson, Stan Boutin, Ben Dantzer, Jeffrey E. Lane, David W. Coltman, Jamie C. Gorrell, Andrew G. McAdam

## Abstract

Organisms can affect one another’s phenotypes when they socially interact. Indirect genetic effects occur when an individual’s phenotype is affected by genes expressed in another individual. These heritable effects can enhance or reduce adaptive potential, thereby accelerating or reversing evolutionary change. Quantifying these social effects is therefore crucial for our understanding of evolution, yet estimates of indirect genetic effects in wild animals are limited to dyadic interactions. We estimated indirect phenotypic and genetic effects, and their covariance with direct effects, for the date of spring breeding in North American red squirrels (*Tamiasciurus hudsonicus*) living in an array of territories of varying spatial proximity. Additionally, we estimated variance parameters and the strength of selection at low and high population densities. Social effects of neighbours on the date of spring breeding were weak at low, but stronger at high population densities. Indirect phenotypic effects accounted for a larger amount of variation in the date of breeding than direct differences among-individuals, although the genetic component to these indirect effects was not statistically significant. Nevertheless, the estimated effect size was large enough to suggest that indirect genetic effects could alter evolutionary change, resulting in less change at high densities despite stronger selection. Despite the difficulty in estimating them precisely, indirect genetic effects have clear potential to alter evolutionary trajectories in any natural systems where organisms interact.

## Introduction

An individual’s phenotype is influenced by both its genotype and the environment it experiences. When organisms live in a social world, mating, competing and cooperating with conspecifics (Frank 2007), the environment they experience is partly made up of the phenotypes of other individuals. This can allow an individual to influence others, and if this trait has a genetic component, then a portion of any organism’s phenotype will be influenced by the genes of those with whom it interacts (Griffing 1967). These are known as indirect genetic effects (“IGEs”), of which maternal genetic effects are a widely known example (Moore et al. 1997; Wolf et al. 1998; McAdam et al. 2014). With maternal genetic effects, the genes of a mother influence the traits of her offspring beyond those directly inherited (e.g. a mammal’s genes affecting milk production may influence the growth rate of her offspring; Koch 1972; María et al. 1993). In the same way, the genes affecting hunting success of an individual (so a direct genetic effect, “DGE”, for condition) in a pack-hunting species may influence the body condition of its pack mates (an IGE). Additionally, when competing for limited resources, the genes for resource acquisition of one individual are expected to negatively influence the resource acquisition, and so resource dependent traits, of individuals with which it competes (Wilson 2014). Therefore, IGEs can be expected in almost any system where conspecifics interact with each other (McAdam et al. 2014).

As IGEs are heritable by definition, they contribute additional heritable variation within a population alongside DGEs (Moore et al. 1997; Bijma and Wade 2008). Similar to a genetic correlation between any two traits (Lande 1979; Kirkpatrick 2009), an individual’s own phenotype for some focal trait and its indirect effect on that trait expressed by neighbours can be genetically correlated (a DGE-IGE correlation). When IGEs are positively correlated with DGEs, which we would expect for the trait of hunting success in the pack-hunter example above, IGEs can enhance that trait’s response to directional selection. This happens because the standard response of the focal trait to selection results in a correlated evolutionary change in the social environment. This in turn causes further change in focal trait mean - in the same direction - through a plastic response to the social environment (Moore et al. 1997).

Conversely, if IGEs are negatively correlated with DGEs then the population response to selection can be reduced, removed, or even reversed (Bijma and Wade 2008; Wilson 2014). Negative correlations are expected for focal traits that are themselves dependent on the outcome of competition for limited resources (Wilson 2014). For instance, Wade (1976) observed a decrease in mean reproductive output across generations in flour beetles (*Tribolium castaneum*) that were under individual selection for *increased* reproduction. This was presumably due to a negative IGE-DGE correlation that caused each subsequent generation to be composed of individuals that more strongly suppressed the reproduction of others through competitive interactions. Similarly, Costa e Silva *et al.* (2013) observed a strong negative DGE-IGE covariance for diameter at breast height in Eucalyptus trees (*Eucalyptus globulus).* This meant that, despite tree growth rates being heritable in the traditional sense (i.e. subject to DGEs), the total heritable variation in the population was near zero, preventing a response to selection. Estimates of DGEs alone might, therefore, provide a poor measure of the potential for a trait to respond to natural selection, yet most estimates of response to selection or evolvability in the wild only consider DGEs (Houle 1992). More specifically, to the extent to which resources are limited in nature, we might expect DGEs to consistently overestimate the adaptive potential of resource-dependent traits because of negatively covarying IGEs (Wilson 2014). As such, IGEs arising from competition represent one possible explanation for the “paradox of stasis”, in which natural selection on heritable traits often leads to stasis rather than evolutionary change (Merilä et al. 2001), yet IGEs are very rarely quantified in the wild.

To date, empirical studies of IGEs in animals have focused on scenarios in which within group interactions can be considered (approximately) uniform, and among-group interactions are absent. This allows IGEs to be estimated from the covariance between phenotypes of group mates, provided pedigree data spanning groups are available (Bijma 2010a). This approach is well suited to dyadic interactions, but also to larger discrete groups (n > 2) of captive animals, where all individuals within a pen are assumed to interact equally, but no interactions occur between individuals in different pens. It has now been applied in a variety of taxa, such as mussels (*Mytilus galloprovincialis*; Brichette et al. 2001), flour beetles (*T. castaneum*; Ellen et al. 2016), Nile tilapia (*Oreochromis niloticus*; Khaw et al. 2016), domesticated chickens (*Gallus domesticus*; Muir 2005; Brinker et al. 2015) mink (*Neovison vison*; Alemu et al. 2014), and domestic rabbits (*Oryctolagus cuniculus*; Piles et al. 2017). This work has helped establish the importance of IGEs for trait evolution (see: Ellen *et al.* 2014, for a review in livestock), and has led to growing interest in studying IGEs in wild populations.

Studies of IGEs in free-living animal populations, however, have thus far been confined to dyadic interactions. For example, Wilson *et al.* (2011) demonstrated that the tendency to win one-on-one fights in wild red deer (*Cervus elaphus*) is subject to both DGEs and IGEs that are perfectly *negatively* correlated, resulting in a total heritable variation of zero. This reconciles quantitative genetic predictions with a common sense approach that sees that the tendency to win cannot evolve at the population level, as each contest must always have one winner and one loser (see also: Wilson et al. 2009; Sartori and Mantovani 2013). Other estimates for IGEs have focused on maternal genetic effects (McAdam and Boutin 2004; McFarlane et al. 2015) or influences of male partner on female bird laying dates (Brommer and Rattiste 2008; Caro et al. 2009; Teplitsky et al. 2010; Liedvogel et al. 2012; Germain et al. 2016). Studies on social interactions in groups of wild animals larger than two are, however, absent.

While social processes in wild populations frequently involve interactions among more than two individuals, it is often problematic to identify and define discrete groups where n > 2. In many cases an individual will interact with multiple conspecifics, but not all at equal intensity. Some interactions are frequent or strong while other interactions are brief or weak (Lusseau et al. 2003; Croft et al. 2004, 2008). One possibility is that organisms interacting in larger groups have generally weaker indirect effects on each of their group mates, as a consequence of their phenotype being “diluted” among more group members (Muir 2005; Hadfield and Wilson 2007; Bijma 2010b). However, within a continuous population (i.e. one in which distinct groups cannot be identified) it seems likely the net effect of one individual on the phenotype of any other may depend on distance or other factors (e.g. time associating) that mediate interaction intensity or frequency (Muir 2005; Cappa and Cantet 2008). To model these situations, variation in interaction strengths can be incorporated as “dilution” or “intensity of competition” factors in IGE models (Muir 2005; Cappa and Cantet 2008; Bijma 2010b). Here we refer to “intensity of association” factors, since social interactions are not always competitive. This approach has proven useful in forestry genetics to estimate DGEs, IGEs, and their covariance, on growth traits in Eucalyptus trees (Costa e Silva et al. 2013, 2017). In this context, the inverse of the distance between each pair of trees was used as the intensity of association factor. The important premise here is that each focal individual has a potential indirect genetic effect on the phenotype of all its social partners, but the degree to which each partner experiences that effect is contingent on its distance from the focal individual. Incorporating intensity of association factors should be equally useful for animal focused IGE models, as allows us to account for animals interacting with multiple different individuals, in groups of varying sizes, and with different individuals at different strengths; a realistic representation of social interactions in the natural world (Fisher and McAdam 2017).

Here we used intensity of association factors to model IGEs amongst multiple neighbours for the first time in a wild animal. We applied this framework to a population of North American red squirrels (*Tamiasciurus hudsonicus,* hereafter “red squirrels”) that have been continuously studied since 1987. We looked at a resource-dependent, but also heritable (h^2^ = 0.14; Lane et al. 2018) life-history trait: parturition date (the date in the spring on which a female squirrel gives birth to a litter; Réale et al. 2003; Boutin et al. 2006; Kerr et al. 2007; Lane et al. 2018), which had the potential to show IGEs. Red squirrels of both sexes in this population live on individual exclusive territories based around a central cache of white spruce (*Picea glauca*) cones called a “midden”. The seeds from stored spruce cones represents their main food source during reproduction in the spring (Fletcher et al. 2013a). Individuals make territorial calls (“rattles”) to delineate territory boundaries (Lair 1990) and deter intruders (Siracusa et al. 2017) from stealing cached resources (Gerhardt 2005; Donald and Boutin 2011). Previous analyses have shown that selection favours earlier parturition dates (Réale et al. 2003), while a food supplementation experiment advanced the timing of spring breeding (Kerr et al. 2007). Note, however, that females can upregulate reproduction prior to a resource pulse (Boutin et al. 2006), and so typically are reproducing below capacity (Boutin et al. 2013), so may not be absolutely food limited. Still, if neighbours compete for food resources, we expect superior competitors to have access to more food and breed earlier. Conversely, competitively inferior individuals are expected to acquire less food and so breed later.

Population density is a key demographic parameter with which we expect IGEs to vary. Selection on birth dates is particularly strong in years of high density (Williams et al. 2014; Fisher et al. 2017; although not found in Dantzer et al. 2013). Furthermore, red squirrels respond behaviourally to both real and perceived increases in density (Dantzer et al. 2012), while mothers adaptively increase the growth rates of their offspring under high density conditions (Dantzer et al. 2013). Taken together, these findings are consistent with the expectation that, all else being equal, high density means increased competition.

In light of the above, we had the following aims and predictions:

1. We expected individuals to have indirect effects on the parturition dates of their neighbours, and that the covariance between direct and indirect effects would be negative. That is, superior competitors will breed earlier and cause their neighbours to breed later (following Costa e Silva et al. 2013; see also: Piles et al. 2017).
2. Parturition dates depend on resource acquisition and possess direct genetic variance, so we expected the indirect effects to possess genetic variance (i.e. to be an IGE; McAdam et al. 2014; Wilson 2014).
3. Increased competition in years of high density should result in stronger indirect effects in high-density years, both in the magnitude of the indirect effects and the strength of the covariance.
4. Finally, we applied the equations of Bijma and Wade (2008) and Costa e Silva et al. (2013) to predict a response to selection in the presence of IGEs, across the whole dataset and at low and high densities separately. This allowed us to assess how IGEs would likely affect evolutionary responses to selection in this system.

## Methods

### Data collection

All data were collected as part of the Kluane Red Squirrel Project in the southwest Yukon, Canada. Since 1987 we have monitored two adjacent and unmanipulated 40 ha. study areas (“Kloo” and “Sulphur”), bisected by the Alaska highway. Red squirrels of both sexes defend exclusive resource-based territories of around 0.3 ha (LaMontagne et al. 2013), centred around a midden, an aggregation of discarded white spruce cone scales underneath which red squirrels cache intact white spruce cones. Each study area is staked at 30m intervals in a grid system and we recorded the x- and y-coordinates of the centre of each midden (to the nearest tenth of a coordinate point, giving distances to the nearest 3m). In the spring of each year we live trapped (Tomahawk Live Trap, Tomahawk, WI, USA) new individuals and gave them unique ear tags in each ear. We also located females (based on vocalizations at known and new territory locations), monitored them for signs of pregnancy and ear tagged their pups once they were born. Based on the previously identified stages of female pregnancy and the body mass of the pups once they were located, we then estimated the female’s parturition date. We analyse this date as the number of days since the 1^st^ January in the calendar year. We also conducted censuses twice yearly (once in spring, once in autumn) using complete enumeration to ascertain the location of all individuals holding a territory, and so estimate population density. See McAdam *et al.* (2007) for further details on the study system.

Red squirrels collect food throughout the summer and autumn, cache it in their middens and rely on it to survive over winter (Fletcher et al. 2013a). The number of cached cones is positively associated with overwinter survival (juveniles: Larivée et al. 2010; juveniles and adults: LaMontagne et al. 2013). Squirrels primarily forage close to their midden, with occasional forays further afield, including small amounts of theft from other red squirrels’ hoards (Donald and Boutin 2011). We define the individuals a red squirrel potentially competes with as its *n* nearest neighbours (*n* was set at 6 for the majority of this analysis, but see below for explorations with different numbers of neighbours). We defined neighbourhoods and population densities based on our autumn census (August) rather than our spring census (May), because autumn is when squirrels are potentially competing for resources to hoard, and conception occurs well before May in most years. Gestation varies little around 35 days (Lair 1985), hence parturition dates cannot be influenced by conditions after conception. Squirrels occasionally defend a second adjacent midden, but as they rarely store food in secondary middens we considered each squirrel’s location to be the location of its primary midden. We then analysed each female’s parturition date the following spring as influenced by her own genes (the DGE), and the identities and genotypes (the IGE) of those competing individuals as identified in the autumn census. Some females gave birth in multiple years, in which case they were included each year they did so, with an updated set of nearest neighbours as necessary. Females may attempt multiple litters in years of high resources, or if their first litter fails (Boutin et al. 2006; McAdam et al. 2007; Williams et al. 2014), but we limited our analyses to each female’s first litter of each year (e.g. Dantzer et al. 2013).

We tagged pups while they were still on their mother’s territory, so maternity is known for all non-immigrants. Male red squirrels provide no parental care. From 2003 onwards, paternities were, therefore, assigned by collecting tissues samples from the ears of adults and neonatal pups. We used these tissue samples to genotype all adults and pups since 2003 at 16 microsatellites (Gunn et al. 2005) analysed with 99% confidence using CERVUS 3.0 (Kalinowski *et al.* 2007; see Lane *et al.* 2007, 2008 for further details). This method gives an estimated error rate of paternities, based on mismatches between known mother-offspring pairs, of around 2% (Lane et al. 2008), which we consider acceptable. Approximately 90% of yearly pups are assigned paternities with known males while the remaining 10% are analysed further in Colony 2.0 (Jones and Wang 2010) to determine whether they might still be full or half siblings from unknown sires using 95% confidence in maximum likelihoods.

### Data analysis

Data on the locations of squirrel territories were available from the autumns in 1991-2015, and so we looked at parturition dates in the following springs (i.e. 1992-2016). All squirrels identified as holding a territory in an autumn census were included in this analysis, including females that did not attempt a litter in the following spring, and males. These individuals had missing values entered for their parturition dates. Their inclusion was nonetheless necessary as they acted as potential competitors during the autumn for those squirrels that did have a litter.

We initially fitted two mixed-effects linear models to our data, the first to estimate indirect effects (the “phenotypic model”), and second to split these indirect effects into genetic and non-genetic components (the “genetic model”). All models we fitted in the software “ASReml” ver 4.1; (Gilmour et al. 2015). We divided raw parturition dates by the standard deviation of all observations, giving a sample with a variance of 1, making the variance components easier to interpret (Schielzeth 2010). In each model we included the fixed effects of study area (a two-level factor), year (as a continuous linear covariate), whether or not the spruce trees “masted” ((produced a super-abundance of cones; Silvertown 1980; Kelly 1994; Lamontagne and Boutin 2007) in the year of the autumn census (a two-level factor), age and age^2^ of the squirrel, and the random effects of year and squirrel identity, to account for repeated measures on squirrels across years. If the age of the squirrel was not known, the mean age of all other squirrels in that breeding season was entered. Estimating the squirrel identity random effect allowed the calculation of the (conditional) repeatability of individual squirrel parturition dates (Nakagawa and Schielzeth 2010). Additionally, while we predicted a negative covariance between neighbours due to competition for resources (especially during high-density conditions), this could be masked by positive spatial autocorrelation in resource availability within a study-area. This would generate a net signal of positive phenotypic covariance among-neighbours (Stopher et al. 2012; Regan et al. 2016; Thomson et al. 2018). To control for this we fitted a term accounting for (non-socially determined) environmental heterogeneity in resource abundance. In our multiyear data set we were unable to obtain convergence from our data with a model in which a separate spatial autocorrelation term for each year was fitted (since the spatial distribution of territory quality is not consistent year-to-year; LaMontagne et al. 2013). As a simpler alternative, we assigned each red squirrel within each year to one 150 m × 150 m square within a grid of non-overlapping squares that encompassed the study area (hereafter referred to as “squares”). Each square was given a unique label comprising its location and the year, and so by fitting this as a random effect we could account for any similarity among red squirrels within each 150 m × 150 m area in each year. This is similar to the approach of Germain et al (2016), who found that an equivalent “grid” term fitted their data better than a matrix of local overlap (c.f. Stopher et al. 2012), or a modelling spatial autocorrelation in the residuals (c.f. Costa e Silva et al. 2013). We repeated this analysis with squares of 75 m × 75 m or 300m × 300 m. These results were qualitatively similar to the analysis with the intermediate size squares, and so are presented in the supplementary materials (Table S1).

To estimate indirect effects, we added the identities of the six nearest squirrels as six random effects (see below for our explorations of other possible neighbourhood sizes), assuming that all random competitor effects came from the same distribution, with a mean of zero and a single variance to be estimated. This allowed us to estimate a single indirect phenotypic effect, and the covariance between this term and the direct effect of squirrel identity. We based “nearest” on location of the primary midden during the autumn census. We associated each neighbour (*j*) of each focal individual (*i*) with variable intensity of association factors (*f*_*ij*_). This allowed the indirect effect of each neighbour *j* actually experienced by *i* to be mediated by their spatial proximity, with *f*_*ij*_ = 1/ (1 + distance), where distance was the Euclidean distance between the center of individuals’ territories, measured in units of 30m. This value is bounded between 0 and 1, with low values representing individuals that were far apart and high values representing individual that were close. We used the inverse of distance here, but any biologically relevant measure representing intensity of social interaction could be used (Fisher and McAdam 2017). To weight the strength of the indirect effects, we replaced all 1s in the indirect effect design matrix with these terms (Muir 2005; Cappa and Cantet 2008). All individuals farther than the 6 nearest neighbours were not modelled as having an indirect effect (but see below). The phenotypic model therefore used the following form, with a population mean accounting for the fixed effects for *i* (*μ*_*Fi*_), a direct phenotypic effect (*P*_*Di*_) and a total indirect influence arising from the sum of competitor specific indirect effects (*P*_*Sj*_) for the 6 nearest neighbours. Note, a single variance for the indirect effect is estimated, from a distribution made up of all competitor effects (see above). Additionally, there are multiple measures per squirrel across years, hence we include the random effect for the year *t* (*K*_*t*_). Our model predicts a parturition date for the ith individual in a given year (*y*_*it*_) and so the residual term is specific to an individual in a year (*e*_*it*_).

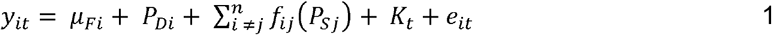

This phenotypic model estimated the variance among squirrels in their parturition dates, the variance in parturition dates caused by repeatable effects of neighbour identities, and the within-individual covariance between direct and indirect phenotypic effects (i.e. *Cov*(*P*_*Di*_, *P*_*Si*_)). For our genetic model, we split these phenotypic effects into additive genetic and permanent environment effects by the incorporation of a pedigree (Kruuk 2004; Wilson et al. 2010). We estimated the DGEs and IGEs on parturition dates, their covariance, and the equivalent terms for the permanent environmental effects:

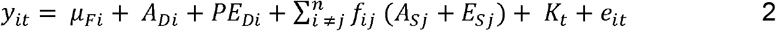

Where individual *i*’s parturition date in year t, is comprised of the fixed effect mean, a direct additive genetic effect (*A*_*Di*_), a direct permanent environmental effect (*PE*_*Di*_), both the additive genetic (*A*_*Sj*_) and non-additive genetic (*E*_*Sj*_) indirect effects of all the *n* neighbours (*j*) that *i* interacts with, a year term (*K*_*t*_), and an individual by year specific residual term (*e*_*it*_).

This model has not been applied to wild animals before, and we fully acknowledge that our choice to consider only the 6 nearest neighbours here is somewhat arbitrary, as indeed is the scaling of *f*_*ij*_. Therefore, we also explored different numbers of neighbours, and different methods for defining our *f*_*ij*_ terms. We then monitored how this influenced the estimates of the variance parameters, to determine whether the model was particularly sensitive to altering these factors (see also: Costa e Silva *et al.* 2017). We present results using *f*_*ij*_ = 1/(1 + distance^2^) in the supplementary materials (Table S1). In the supplementary materials we also present results where we defined the competitors as all those within 60, 130 or 200 metres, without weighting by distance, up to 24 competitors (Table S1), and investigations with varying numbers of neighbours 1-5, 9, 12, 15, 18 & 24; Table S2). Neither changing the number of neighbours nor rescaling intensity of association terms changed the number of model parameters estimated. Therefore, information criteria-based approaches for comparing model fits were not appropriate. Additionally, we were primarily interested in our ability to estimate, and the magnitude and significance of, certain parameters (our indirect effects), hence finding the most parsimonious model of parturition date was not a goal of ours. Instead we simply assessed the change in variance components, noting the size of the parameter estimates and size of the standard errors. We focus on the results with the 6 closest neighbours, as this seemed the median result among the variations we tried. Using the inverse of distance^2^ squared led to a large increase in the standard errors of the DGE estimate, which only occurred in this model, hence we considered simply the inverse of distance as more appropriate.

We tested the significance of the direct-indirect phenotypic covariance in the phenotypic model using a likelihood ratio-test (LRT) between a model with the covariance freely estimated and one with it fixed to zero, and tested the significance of the indirect phenotypic effect using a LRT between the model with the indirect effect (and a zero covariance) and a model without it. With the genetic model, we tested the significance of the DGE-IGE covariance, and the IGE variance, in the same way, in models that still estimated the full direct-indirect phenotypic covariance matrix. We assumed the LRT statistic was distributed as a 50:50 mixture of χ^2^_1_ and X^2^_0_ when testing single variance components (following Self and Liang 1987) but as X^2^_1_ when testing covariances. Note that, even where terms were non-significant, they were retained as our best estimate of the corresponding parameter for our estimates of predicted responses to selection (as described below). We report correlations, although if the variance of either the direct or indirect effect was very small (<0.0001), then we assumed it was essentially zero, and so then we report the correlation as “undefined”. Although they were not directly relevant to the biological hypotheses being tested, the statistical significance of the fixed effects in the genetic model was tested using conditional Wald tests (see: Gilmour et al. 2015). This approach to testing the significance of fixed effects in mixed linear models performs well in situations with limited sample sizes (Kenward and Roger 1997). We then calculated partial R^2^ for each fixed effect, following Edwards et al. (2008), using the residual degrees of freedom as calculated by ASReml (1174 for the genetic model).

### Influence of population density on indirect effects

We consider population density during the resource caching period to be key to resource acquisition. Consequently, for any given year of parturition the relevant measure of density was obtained from the census in the autumn of the year *prior* to parturition, i.e. at the same time as when the territory ownership was defined. As the study area has grown marginally since the start of the project, we restricted counts to individuals holding a territory within a defined 38ha area that has been constant throughout the entire study period. Across both study areas in all years the median population density in the autumn was 1.69 squirrels ha^−1^ (Fig. 1). We, therefore, labelled each study area within each year with a density higher than this as “high density” (1994, 1998-2000, 2006 and 2015 for both study areas, 1991-1993, 1995-1997, 2001 and 2002 for Sulphur only and 2011-2014 for Kloo only), and so the remainder as “low density” There were, therefore, 26 instances of low density conditions, and 24 instances of high density conditions. There are several instances of study areas having exactly the median density, hence why there are more low-than high-density conditions.

**Figure 1.**
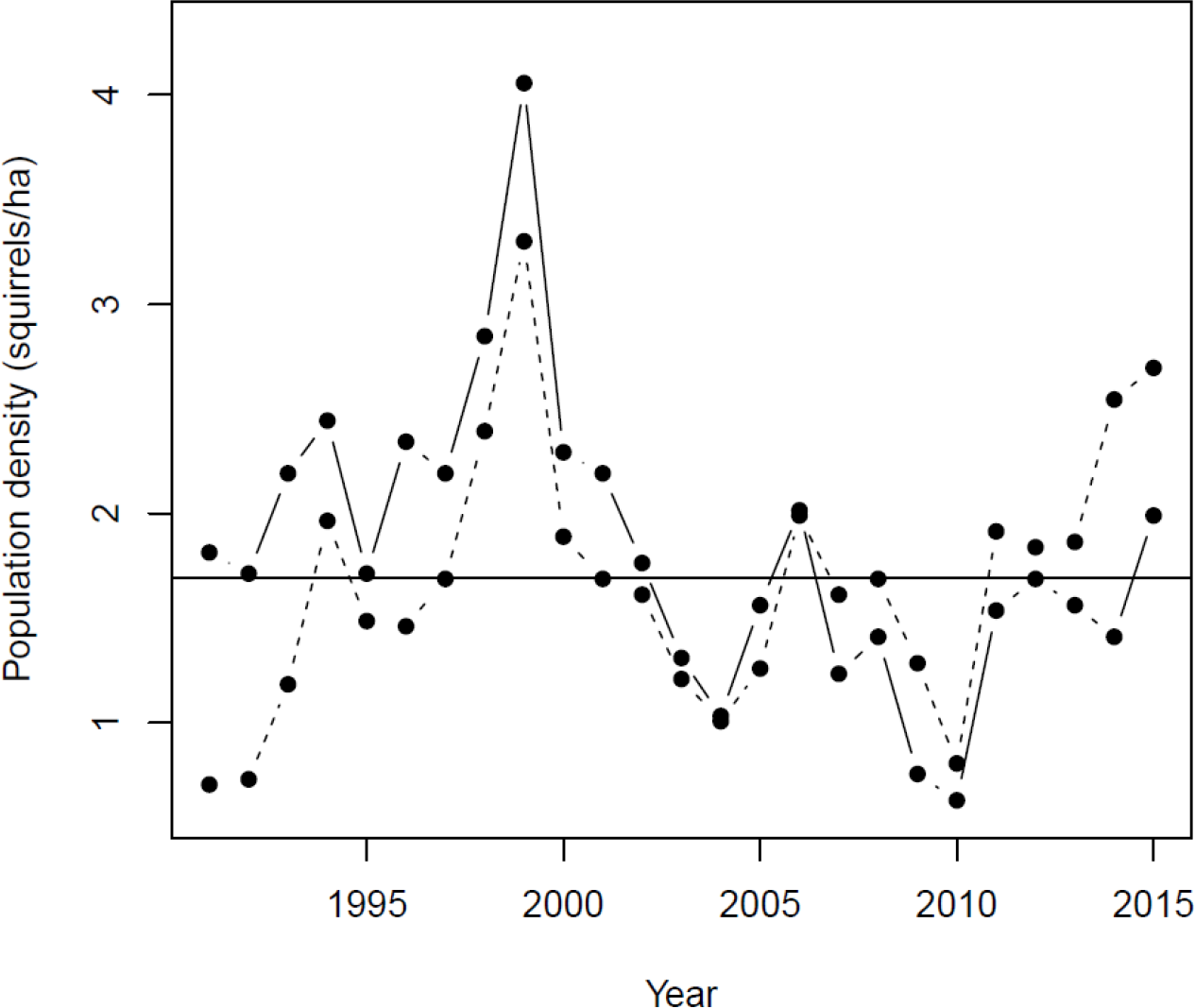
Estimated population densities across both study areas in our study (“Kloo” is solid line, “Sulphur” is dashed line). Points above the line (the median density: 1.69 squirrels ha.^−1^) were counted as “high density”, points below the line as “low density”.

**Figure 2.**
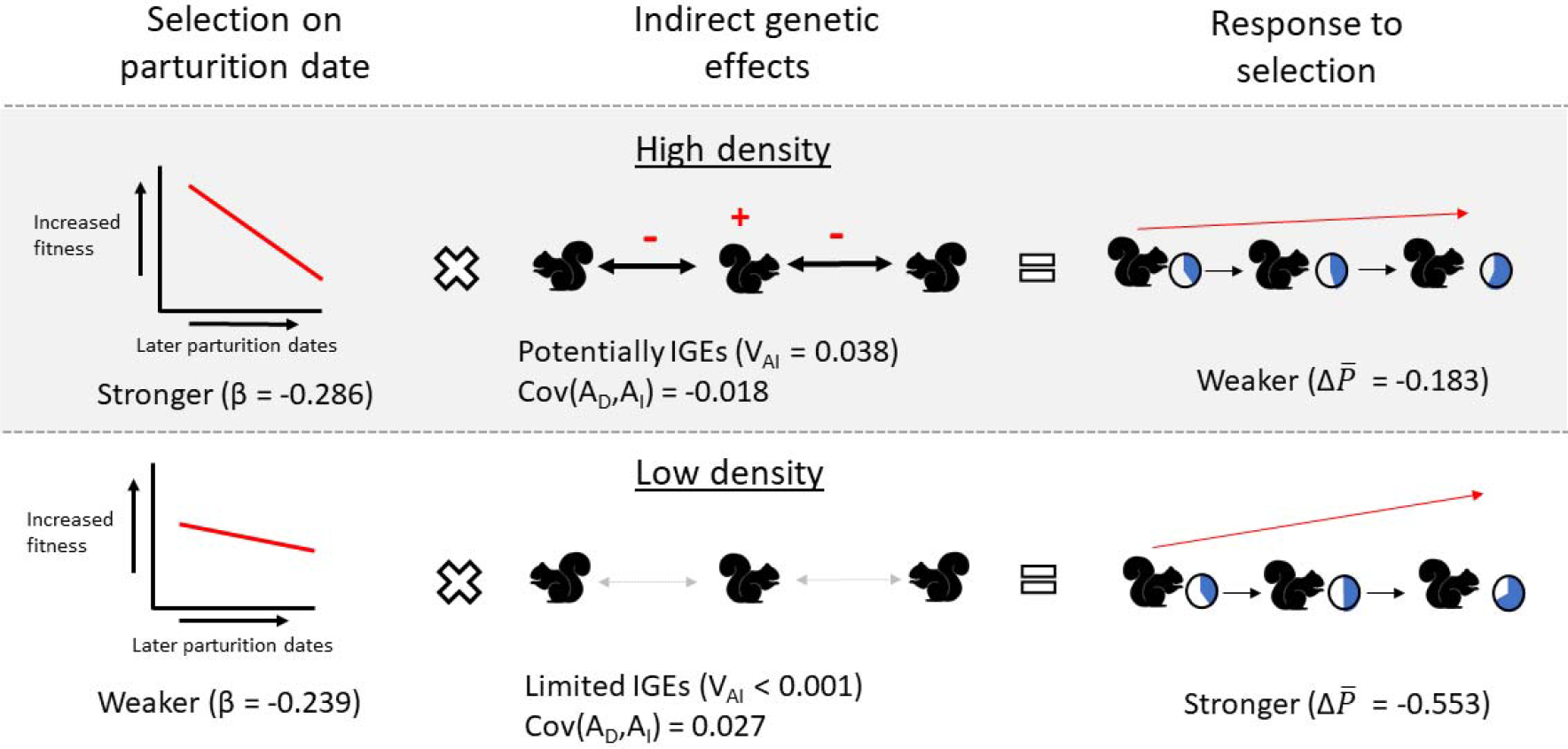
A summary of our results for the analysis of indirect genetic effects at low and high densities. At high density selection was stronger (β = −0.286), and there were direct genetic effects (variance = 0.053). There were significant phenotypic indirect effects, and non-zero but statistically non-significant indirect genetic effects (variance = 0.038). The DGE-IGE covariance (Cov(A_D_,A_I_)) was estimated to be negative (estimate = −0.018). These parameters gave a weaker response to selection (Δ*P̅* = −0.183 days.generation^−1^). At low density selection was weaker (β = −0.239), while there were direct genetic effects of a similar magnitude to at high densities (variance = 0.047), but indirect genetic effects were absent (variance = < 0.001). The DGE-IGE covariance was estimated still (estimate = 0.027). These parameters gave a moderate response to selection (Δ*P̅* = −0.553 days.generation^−1^). Note in all cases negative A*P̅* values indicate the evolution of earlier parturition dates. For the response to selection, more full circles represent later parturition dates. Red symbols represent the DGE-IGE covariance, showing that individuals with genes to be early (red plus) cause their neighbours to give birth later (red minus).

For both the phenotypic and the genetic models, we fitted an interaction between population density (low or high) and each random effect. This gave us separate density-specific estimates of each of the variances (DGEs, IGES, and non-genetic versions) and covariances, the among-year variances and the among-square variances for low- and high-density study areas. To obtain stable model convergence in the genetic model, we were required to fix the direct permanent environment effect in low-density years to 0.1 × 10^−4^, but since this term was estimated to be very small in the model across all years, this is likely not problematic. There was a single residual variance in each model. We also included density as two-level factor in the fixed effects, and an interaction between this term and each of the other fixed effects, to allow them to vary between low-high-density conditions. We tested for significance of indirect effects in both low- and high-density conditions in the same way as for the full models. When testing the significance of terms for low-density, we maintained the full structure (e.g. IGEs and their covariance with DGEs in the genetic model) for high-density conditions, and vice versa.

### Calculating variance parameters and the predicted response to selection

To assess the contribution of indirect effects to phenotypic and genetic variances, we estimated the variance in individuals’ phenotypic effects on the population mean parturition date (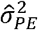, incorporating both consistent direct and indirect phenotypic effects; for the phenotypic model), and variance in individuals’ heritable influence on the population mean parturition date (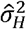; for the genetic model, commonly referred to as the “total heritable variance”). Following Bijma (2011) and Costa e Silva *et al.* (2013) these are:

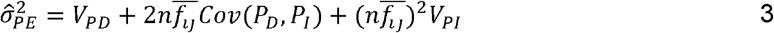

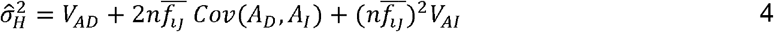

Where *n* is the number of neighbours (excluding the focal individual, so 6), 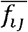 is the mean intensity of association factor, *V*_*PD*_ and *V*_*AD*_ are the direct phenotypic and additive genetic variances respectively, *Cov*(*P*_*D*_,*P*_*I*_) and *Cov*(*A*_*D*_,*A*_*I*_) are the phenotypic and genetic direct-indirect covariances respectively, and *V*_*PI*_ and *V*_*AI*_ are the indirect phenotypic and additive genetic variances respectively. The 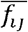 was calculated as 0.330 across the whole dataset, 0.298 at low densities and 0.352 at high densities, which means a squirrel’s 6 nearest neighbours were on average, 60.9m, 70.7m and 55.2m from it across the whole dataset, at low densities, or at high densities respectively. Note that 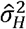, unlike traditional heritability, can exceed 1; see Bijma (2011) for the mathematical demonstration of this, and Ellen *et al.* (2014) for empirical examples in livestock.

In order to calculate the predicted response to selection in the presence of IGEs among related individuals, we combined equation 15 of Bijma and Wade (2008), and equation S5_2 of Costa e Silva et al. (2013), setting total phenotypic variance as 1. In a previous study (Fisher et al. 2017) we determined that selection on parturition date acts primarily among neighbours within 130m, and, so the relative strength of multilevel selection (*g*) is approximately zero, giving:

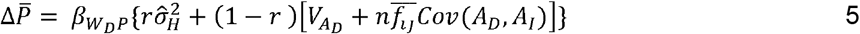

Note that as we have already defined *n* as the number of neighbours excluding the focal individual, we have altered *n-1* in eq. 15 of Bijma and Wade (2008) to *n*.

In eq. 5 the change in the mean phenotype (Δ*P*̅) is predicted by the selection gradient of the phenotype on relative direct fitness 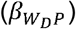, multiplied by a term encompassing the genetic variance parameters. We estimated 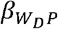 across all females in the whole dataset by regressing relative fitness (number of pups born to each individual in a given year that recruited to the population as adults, divided by the population average for that year) on standardised parturition dates (mean centred and divided by the standard deviation across the whole dataset). This measure of fitness maps individuals that recruited into our population (and hence held territories, necessary to be included in our analysis) to the number of recruits they have. Arguably, early survival is a trait of the juvenile, not the parent, and hence this fitness component should not be assigned to the parent (Thomson and Hadfield 2017). However, as parturition dates are only expressed by females, and do not influence adult survival (Lane et al. unpublished), our estimates of selection should not be biased. We re-estimated selection gradients at low and high densities by fitting an interaction with low/high density to parturition date, to give different estimates for 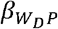 in each condition. Densities (and competitive neighbourhoods) in the autumn of a given year were associated with parturition dates and fitness in the following spring (year +1). Relatedness (*r*) is the mean coefficient of relatedness among the 7 squirrels in each neighbourhood (the focal squirrel and its 6 nearest neighbours). We used the pedigree to calculate *r* as 0.094 across the entire data set, 0.087 at low densities and 0.098 at high densities. We multiplied Δ*P̅* by the standard deviation of parturition date, to give the result as “days per generation”. We compare this to a case where IGEs were equal to zero, and so the response to selection was equal to 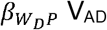.

## Results

In total, 1862 unique red squirrels were recorded a total of 4362 times in autumn censuses as holding territories, and so were included in the analysis. There were 555 unique females that had at least one litter, with a mean of 2.1 (range = 1-8, standard deviation = 1.3) recorded parturition dates each. The median date of first litters was 23rd April, with interquartile ranges of 6th April to 11th May. There were 364 females that had no recorded parturition dates, and 943 males. 1196 squirrels had a known mother, and 498 had a known father, with 481 of those having both parents known.

Parturition dates differed greatly among years and less so among squares, with variance among-years accounting for 32.0% of the variance in the genetic model, while variance among-squares accounted for 4.0% of the total variance (all variance component estimates are shown in Table 1, with fixed effect estimates shown in Table 2). While there was no linear trend across years, parturition dates were significantly earlier following mast years by approximately 40 days.

Alongside these environmental variations, individuals showed some degree of consistency in their parturition dates, with the direct variance among-individuals in parturition date in the phenotypic model accounting for 3.8% of the phenotypic variance. Indirect phenotypic effects of neighbours were significant (LRT, X^2^_0.5_ = 13.755, p < 0.001), but the covariance between the direct and indirect phenotypic effects was not (cor = −0.094, LRT, X^2^_1_ = 0.111, p = 0.739), indicating that individuals that give birth earlier do not influence their neighbours in any particular direction relative to their own parturition date. Individuals’ consistent differences in their own phenotypes and consistent effects on neighbours 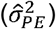 was calculated as 31.4% of the phenotypic variation, indicating that social effects account for a large amount of the variation in parturition date. Alongside this consistency, individuals showed a degree of plasticity, with older squirrels having earlier parturition dates, while the positive quadratic effect indicates a nonlinear effect of age in which squirrels began to breed later at older ages.

Parturition date showed direct heritability, with V_AD_ in the genetic model accounting for 4.8% of the phenotypic variance (note this differs from previous estimates of *h*^2^ for this trait in this system as here we include the among-year variation in *V*_*P*_). The estimate for the IGEs was not different from zero (LRT, X^2^_0.5_ = 0.003, p = 0.480), as was the DGE-IGE covariance (cor = undefined, LRT, X^2^_0.5_ = 0.1 19, p = 0.729). We calculated the total heritable variance of parturition date, 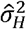, as 6.8%, a modest increase over V_AD_. The very small DGE-IGE covariance indicates that genotypes for early parturition dates did not affect their neighbours in any consistent direction relative to their own parturition date.

### Low vs high density comparison

In low density conditions, both the variance in indirect phenotypic effects (LRT, X^2^_0.5_ = 0.808, p = 0.184) and the direct-indirect phenotypic covariance (cor = 0.737, LRT, X^2^_0.5_ = 0.1.206, p = 0.272) were not significantly different from zero. At high densities there were significant phenotypic indirect effects (Table 1; LRT, X^2^_0.5_= 9.523, p = 0.001), although the covariance was not different from zero (cor = −0.023, LRT, X^2^_0.5_ = 0.004, p = 0.952).

Given that we detected no phenotypic indirect effects in low-density conditions, it is unsurprising that the IGEs (LRT, X^2^_0.5_ = 0.000, p = 0.500) and the DGE-IGE covariance in these conditions were also not different from zero (cor = undefined, LRT, X^2^_1_ = 0.566, p = 0.452). For high densities, IGEs were considerably stronger than across the whole dataset, and more than one standard error from zero, although still not significantly different from zero (LRT, X^2^_0.5_ = 0.607, p = 0.218). The covariance between DGEs and IGEs was negative but not different from zero (cor = −0.401, LRT, X^2^_0.5_ = 0.688, p = 0.407). Although we reiterate that neither covariance was statistically significant, based on our parameter estimates in low-density conditions 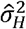 was 14.3%, which was higher than V_AD_, as this calculation includes the positive DGE-IGE covariance estimate (despite the lack of variance in IGEs rendering the correlation undefined). In high-density conditions 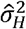 was 14.2%, much higher than with direct genetic effects alone due to the additional genetic variance from IGEs. We stress that, as the estimates for the IGEs and their covariances with the DGEs were not significantly different from zero, the estimates of 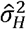 should be interpreted with caution.

The square term revealed that there was some variation attributable to spatial location in both conditions, accounting for 4.2% in low-density, and 3.1% in high-density conditions, of the variance in the genetic model split between low and high densities. Finally, there was also substantial among-year variance in both conditions, accounting for 32.2% and 38.4% for the observed variance in low and high-density conditions respectively. We present estimates for fixed effects at low and high densities from the genetic model in the supplemental materials (Table S3); for the calculation of partial R^2^s, we calculated the residual degrees of freedom to be 1169.

### Predicted response to selection

Across all years, selection favoured squirrels with earlier parturition dates (linear selection gradient β = −0.249). The standard deviation of parturition date was 23.32. From eq. 5, this gives a predicted response when accounting for IGEs across the entire data set of −0.342 days generation^−1^, or −0.279 days generation^−1^ if IGEs were not considered. While the magnitude of selection differed, breeding earlier was still favoured whether squirrels were breeding under low (β = −0.239) or high densities (β = −0.286). At high densities, despite stronger selection, we calculated a slower predicted evolutionary response, due to the negative DGE-IGE covariance. At low densities, updating all parameters except *n*, the predicted response was −0.553 days generation^−1^, while at high densities it was −0.183 days generation^−1^. Predictions solely based on additive genetic variance multiplied by the selection gradient were −0.262, and −0.353 days generation^−1^, for low densities and high densities respectively, therefore giving a faster response when selection is strongest, as is typical.

## Discussion

### Indirect effects are present and change with population density

Red squirrels live in territories surrounded by conspecifics, with whom they engage in social interactions through vocalizations, competition for resources, and mating interactions. Our analyses show that these interactions can lead to substantial indirect effects on female squirrel reproductive traits. These are detected here as a repeatable influence of competitor identity on the parturition date of focal individuals - which accounted for a much greater amount of variation in parturition date than direct effects of individual identity alone. Our results also suggest that these indirect effects are more important determinants of focal phenotypes at high densities than at low densities. Specifically, at high densities, there is significant variation in the extent to which squirrels influence each other’s parturition dates, but this is not the case at low densities.

**Table 1.**
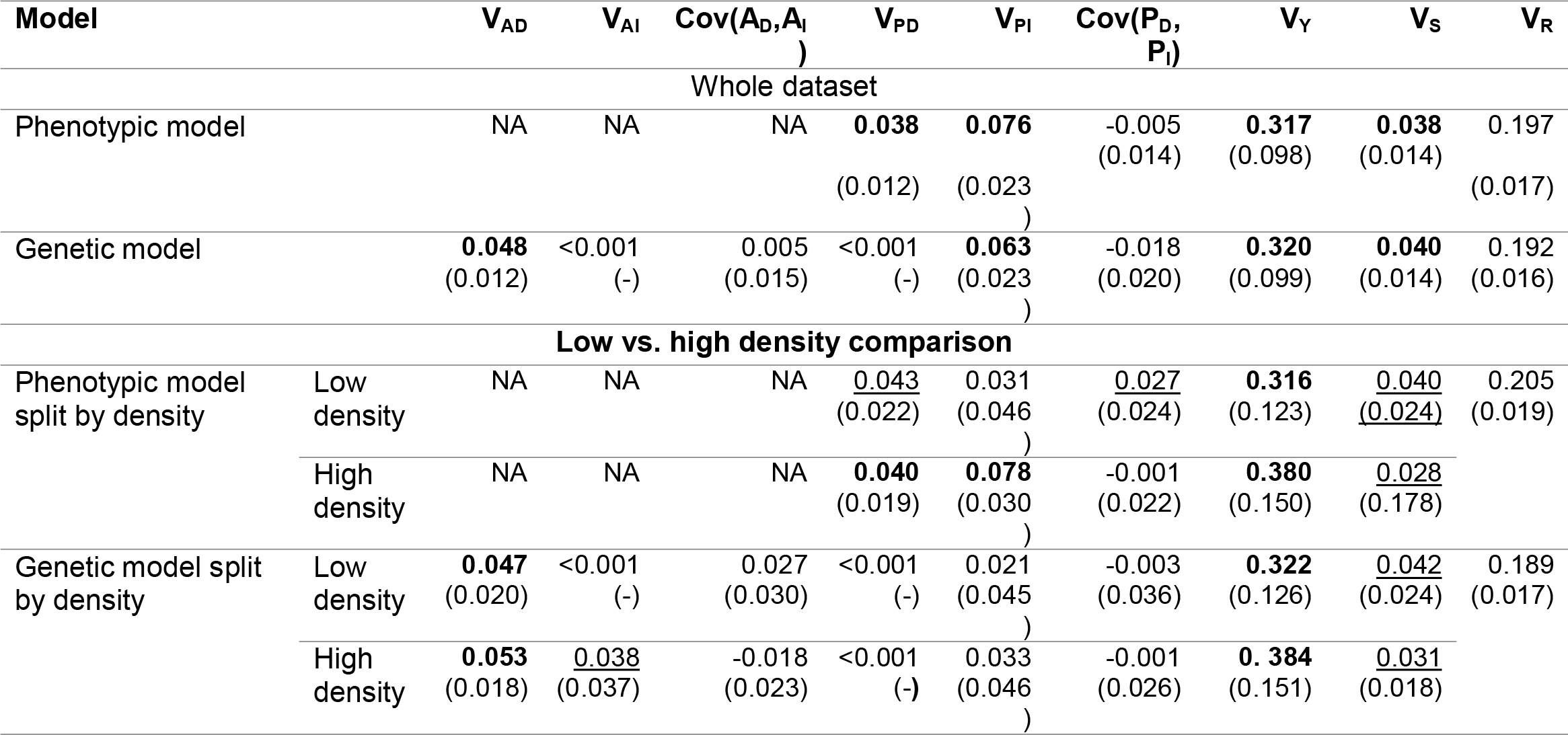
Variance component estimates (with their approximate standard errors in brackets) for each element of the variance-covariance structure from our models. Terms that were bound to values very close to zero will not have a standard error estimated, and so have “-“ instead. Models without a given term have “NA” entered in that cell. Terms highlighted in bold were >2 times greater than their standard errors, while terms underlined were between 1 and 2 times greater than their standard errors. Variance in direct genetic effects are indicated by V_AD_, in indirect genetic effects by V_AI_, and their covariance by Cov(A_D_, A_I_). Equivalent notation with “P” instead of “A” refers to variance in purely phenotypic effects for the phenotypic model, and permanent environment effects in the genetic model. V_s_ is the among-square variance (with squares of size 150m×150m), V_Y_ is the among-year variance, and V_R_ is the residual variance.

**Table 2.**
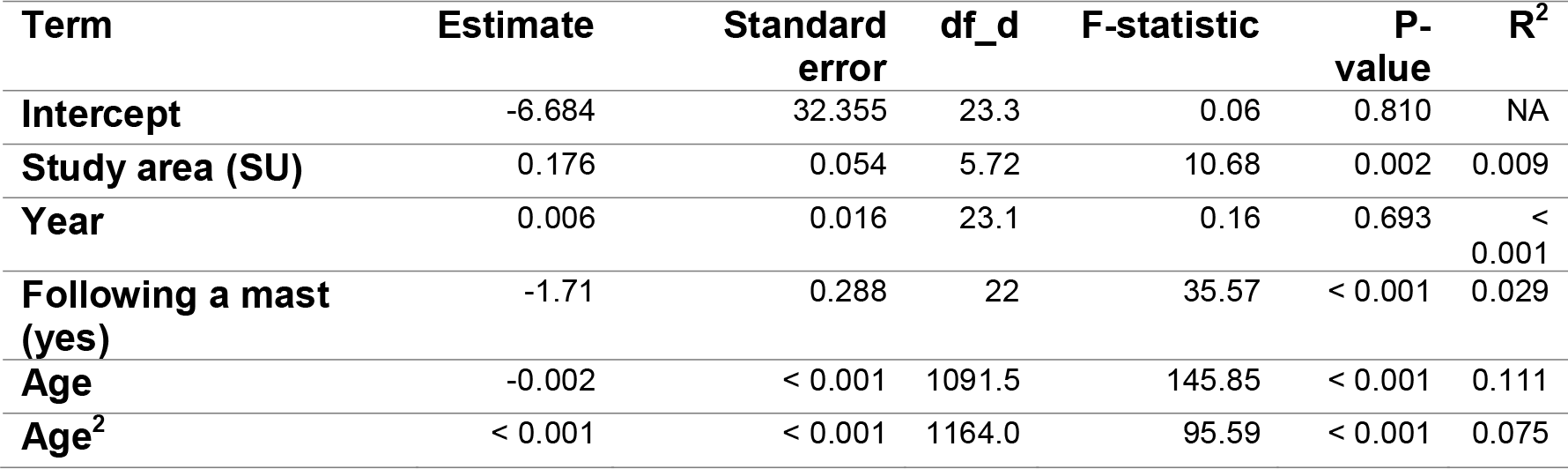
Estimates and relevant statistics for fixed effects from final model with all years. Study area was a two-level factor, with “Kloo” as the reference level, hence the shown estimate is for the deviation of the “Sulphur” study area. Following a mast was a two-level factor, with not following a mast as the default, hence the estimate is for the deviation in parturition dates following a mast year. The denominator degrees of freedom are indicated by df_d, while the numerator degrees of freedom were 1 in all cases.

| Term | Estimate | Standard error | df_d | F-statistic | P-value | R<sup>2</sup> |
| --- | --- | --- | --- | --- | --- | --- |
| Intercept | -6.684 | 32.355 | 23.3 | 0.06 | 0.810 | NA |
| Study area (SU) | 0.176 | 0.054 | 5.72 | 10.68 | 0.002 | 0.009 |
| Year | 0.006 | 0.016 | 23.1 | 0.16 | 0.693 | < 0.001 |
| Following a mast (yes) | -1.71 | 0.288 | 22 | 35.57 | < 0.001 | 0.029 |
| Age | -0.002 | < 0.001 | 1091.5 | 145.85 | < 0.001 | 0.111 |
| Age<sup>2</sup> | < 0.001 | < 0.001 | 1164.0 | 95.59 | < 0.001 | 0.075 |

The social effects on parturition date we documented indicate that much more of an individual’s phenotype is under the control of those it socially interacts with than is determined by its own identity, even in a solitary and territorial species. Work on Eucalyptus trees (Costa e Silva et al. 2013) implicated competition for limited resources as the source of indirect effects, and our results are consistent with this idea, hence highly competitive red squirrels may acquire larger amounts of resources from the environment, leaving less for other individuals. Earlier studies have shown that red squirrel females may be food limited to some degree, aside from in years following a mast event. For example, earlier parturition dates and lower levels of oxidative protein damage and higher levels of antioxidants were found when food was supplemented (Kerr et al. 2007; Fletcher et al. 2013b; Williams et al. 2014), and individuals are more likely to survive over winter with a larger food cache (Larivée et al. 2010; LaMontagne et al. 2013), suggesting that not all individuals have enough stored food. However, female squirrels appear to reproduce below capacity in non-mast years, and upregulate their reproduction *before* pulsed resources are available (Boutin et al. 2006, 2013), and so they are likely not completely food-limited. The additional insight from the current study is that, for focal individuals, competitive effects on phenotype depend not simply on high density, but also on the identities - and so phenotypes - of their nearest neighbours.

Our analysis did not explore the specific mechanism (or trait(s)) that mediate indirect phenotypic effects from competition, hence we have not confirmed that red squirrels are competing for limited food resources, although this explanation seems likely. We can however, suggest a second explanation based on prior knowledge of the system: red squirrels might influence each other’s parturition dates through acoustic territorial interactions. Red squirrels give territorial calls (“rattles”), to which neighbours behaviourally respond (Shonfield 2010) and which function to maintain their territory from conspecifics (Smith 1978; Lair 1990; Siracusa et al. 2017). Additionally, red squirrels rattle more when they have a higher local population density (Dantzer et al., 2012; Shonfield et al. 2012), while red squirrel mothers increase the growth rate of their pups when playback of territorial vocalizations leads to the perception of higher local population density (Dantzer et al. 2013). This is through upregulation of maternal glucocorticoids, part of the stress axis (Dantzer et al. 2013). Other life history traits, such as parturition date, may be influenced by rattles at high densities, allowing individuals to influence each other’s parturition dates. Therefore, acoustic interactions among-neighbours, which enable neighbours to influence each other’s reproduction, may be a source of indirect effects, particularly in high-density conditions.

### Indirect effects with a limited heritable basis

While our analyses provide statistical support for considerable indirect effects of competitors on a focal individual’s parturition date, we were unable to conclusively demonstrate that these indirect effects were underpinned by genetic variation. Estimated effect sizes were larger at high densities, in line with our predictions and the phenotypic effects. If we incorporated the estimates of variance attributable to IGEs, and the DGE-IGE covariances, into our predicted response to selection, we expected a faster response to selection in low-density conditions (−0.553 days generation^−1^) when selection was weakest, while a slower response to selection in high-density conditions (−0.183 days generation^−1^) when selection was strongest, compared to across the whole data set (−0.342 days generation^−1^). Therefore, a negative DGE-IGE covariance counteracted stronger selection at high densities to give a slower evolutionary response. However, while the point estimates of predicted change indicate IGEs are potentially strong enough to make a meaningful difference to evolutionary dynamics, we acknowledge they are estimated with high uncertainty.

Previous work on livestock has shown that IGEs negatively correlated with DGEs can reduce or even reverse the expected response to selection (Costa e Silva et al. 2013; Muir et al. 2013; Ellen et al. 2014), as we have found. The evolutionary stasis of heritable traits under directional selection is a well-known observation in need of an explanation in the study of trait evolution in wild populations (Merilä et al. 2001; Kokko et al. 2017). IGEs that consistently counteract selection responses (compared to a DGE-only scenario) would reduce evolutionary change, as we have shown under high-densities, and so could contribute to a lack of evolutionary change. Whether this is a general explanation for evolutionary stasis remains to be explored (Wilson 2014). In our study population, despite phenotypic selection on parturition dates (which as noted above are heritable), we have observed no evolution in this trait over 20 years (Lane et al. 2018). However, Lane et al (2018) found that the association between parturition date and fitness was entirely a residual correlation, rather than a genetic one, so no alternative explanation for evolutionary stasis (such as IGEs) is required.

The contribution of the variance of the IGEs, and the DGE-IGE covariance to the expected response to selection is not certain, as these estimates were not statistically significant. If IGEs are not different from zero, then the expected response to selection may not differ from that predicted by the breeder’s equation (Bijma and Wade 2008). We note that the non-significance of our IGE variance estimates may have been driven by a high degree of uncertainty (large standard errors), rather than the magnitude of the effect, as in high density years the V_AI_ was quite close in absolute size to V_AD_, and their contribution to total heritable variance and the predicted response to selection was large enough to change the expected response in different conditions. As such, the value of incorporating these estimates into predictive models is possibly large, but uncertain.

Predictions about the speed of evolution based on *V_AD_* or 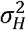 can be misleading, because, as we have demonstrated, a negative DGE-IGE covariance detracts from the response to selection (see eq. 5), even if there is a large amount of genetic variance. This is similar to how negative genetic correlations between traits under equivalent selection can limit their evolution (Lande 1979; Kirkpatrick and Lande 1989). Additionally, there are several reasons why not all of 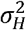 may be utilized in the response to selection. As can be seen from eq. 5, the response to selection in the presence of IGEs is not simply 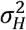 multiplied by selection. Neighbourhoods or groups made up of unrelated individuals, small group sizes, and a near-zero or negative covariance between DGEs and IGEs will also cause the 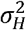 to be somewhat discounted when calculating the response. However, given their prevalence among livestock (Ellen et al. 2014), IGEs may well influence the response to selection more broadly, hence they need to be considered more often when attempting to predict the microevolution of populations, and to explain phenomena such as evolutionary stasis (Merilä et al. 2001; Ellen et al. 2014; McAdam et al. 2014). This is true even in populations of wild animals that do not live in clearly defined groups, but in an irregular network of territories (see also Nunney (1985) for related work on the evolution of altruism in “continuous arrays” of animals).

### Altering competition indices and neighbourhood size

Varying the intensity of association factors (i.e. how strongly we weighted neighbours at different distances) and the size of the neighbourhood did alter the balance between the estimated direct and indirect effects, as well as estimated relative contribution of genetic and environmental influences (see Tables S1-2 in the supplementary materials). Weighting the closest individuals more strongly, by only including the 1-3 nearest neighbours, or using the inverse of distance or distance^2^, or by only including individuals within 60 m, gave similar results. In all these versions, the variance arising from DGEs increased marginally compared to the model where all neighbours were weighted equally. This effect was more pronounced when using the inverse of distance^2^ to define the intensity of association factors. We note that the standard errors of estimates for direct additive genetic variance (V_AD_) in the model using the inverse of distance^2^ were greatly increased, causing the estimate to be within two standard errors of zero (i.e. nominally non-significant). This was the only model explored where this occurred. Weighting farther individuals as strongly as close individuals, either by not including any intensity of association factors for the 6 closest individuals, or by including all individuals within 200 m and weighting them equally, gave very low estimates for the IGEs. This could suggest that individuals at greater distances do not influence their neighbours as much as close individuals.

Increasing the number of neighbours considered in the analysis beyond six led to larger estimates for the variance arising from the non-genetic indirect effects (V_PI_). A larger estimate for the V_PI_ was also present in the model before the square term was added (not shown). This suggests the apparent non-genetic influence of neighbours at large spatial scales, as indicated by V_PI_, may be driven by shared environmental factors at the larger scale causing sets of neighbours to be consistently different from other sets, rather than by social interactions of the focal individual causing their neighbours to be consistently different. Decreasing the number of neighbours tended to increase the variance attributed to the DGE, while IGEs showed a non-linear trend, peaking in magnitude with 4 neighbours and then falling back down towards zero. At these neighbourhood sizes, V_PI_ was typically estimated near zero, but grew in size once 5 or more neighbours were considered. Overall, these results do not indicate that inferences from our model with the six closest neighbours, weighted by the inverse of distance, are inappropriate for the system.

The approach we used, based on the work of Muir (2005) and Cappa and Cantet (2008) can be applied to organisms in a range of social structures. Due to the relatively recent increase in usage of techniques such as social network analysis (Krause et al. 2007, 2014; Croft et al. 2008), estimates of pairwise associations within populations of animals have been made in many systems. These values can be used as the intensity of association factors, as we used the inverse of distance, to scale indirect effects (Fisher and McAdam 2017). To estimate IGEs, this must be twinned with information on the phenotypes and relatedness of the individuals in the population. We had a large dataset with good information on phenotypes and relatedness of individuals, yet high uncertainty around moderately large estimates of IGEs did not distinguish them from zero. The requirement to phenotype, genotype and assess the social relationships of many individuals within a population may well limit the range of study systems this approach can be used in (Kruuk and Wilson 2018). However, with decreases in the cost of tracking technologies and in the cost of assessing the genetic relatedness of animals (Bérénos et al. 2014), more study systems will begin to be able to apply this and similar models, increasing the number of estimates for these difficult-to-estimate quantitative genetic parameters, which could then be aggregated in a meta-analysis to detect general patterns (Reid 2012), such as that by Wilson and Réale (2005) for the direct-maternal genetic correlation.

### Conclusions

Previous to this study, IGEs had only ever been estimated for wild animals in the context of pairwise (dyadic) social interactions. We extended this to estimate IGEs on a life-history trait with links to fitness in a population of wild animals that do not interact in discretely defined groups. We also incorporated varying strengths of closeness of association between individuals to more accurately represent the heterogeneous and complex nature of social interactions in the natural world. We found that indirect effects of neighbours were a very important contributor to parturition dates, especially at high densities, and may have a heritable component. Predicting selection responses from a model that incorporated IGEs indicates that they can both slow down (at high population density) and speed up (at low density) the expected response to selection for earlier parturition dates. This is despite selection actually being stronger at high densities. However, these patterns are based on point estimates for genetic parameters that are characterised by high uncertainty and, as noted, we cannot exclude the possibility that the indirect effects have a non-genetic basis. Nonetheless, significant indirect phenotypic effects were detected and appear to increase in importance at high density. This is consistent with competition for limited food resources being the source of neighbour influences on focal life-history traits. Exactly how this competition is mediated remains to be determined. The estimation of indirect effects, and IGEs specifically, should be extended to more systems where densities and resource availabilities vary (either naturally or artificially) to determine whether the patterns we have observed are general. While we did not conclusively demonstrate IGEs are present, we think wider estimation of effect sizes is useful even if power is limiting to make strong inferences in any single case. The method we have used is flexible enough to be applied to alternative systems, hence we look forward to the accumulation of more estimates of IGEs in the wild to detect general patterns.

## Acknowledgements

We thank Agnes MacDonald for long-term access to her trapline, and to the Champagne and Aishihik First Nations for allowing us to conduct work on their land. We thank all the volunteers, field assistants and graduate students whose tireless work makes the KRSP possible. We thank Bill Szkotnicki and Piter Bijma and assistance with the analyses. We have no conflicts of interest.

## Author contributions

AGM, AJW, and DNF conceived of the research question. SB initiated the long-term study and all authors contributed to field logistics, data collection and the writing of the manuscript. DNF drafted the manuscript and conducted the data analysis, with guidance from AJW and AGM. All authors approved of the final manuscript for submission.

